# Midgestation metabolic constraint in purine metabolism drives distinct strategies for placenta and fetal growth

**DOI:** 10.64898/2026.03.18.712680

**Authors:** Weizhi Xu, Nancy De La Cruz, Andrea Woods, Dmitry Lokshtanov, Shihong Gao, Nawal Khan, Sylvia Wright, Maria E. Florian-Rodriguez, Donald D. McIntire, Elaine L Duryea, David B. Nelson, Catherine Y. Spong, Christina L. Herrera, Jacob H. Hanna, Sanjay Srivatsan, Alejandro Aguilera-Castrejon, Ashley Solmonson

## Abstract

Purine nucleotides are essential for mammalian development^1,2^. Purine monophosphates support cell signaling and proliferation and are synthesized by cells through either de novo synthesis or a salvage pathway^3^. We previously identified a midgestational metabolic transition in mice (gestational days gd10.5–11.5) characterized by changes in purine metabolism^4^. Midgestation is a period of rapid growth for placenta and embryo, yet it remains unclear how the placental tissues expand without directly competing with the embryo for biosynthetic resources. Here, we show that this midgestational metabolic transition is associated with a marked reduction in embryonic expression of purine salvage enzymes, which constrains embryonic metabolism and leads to different strategies for purine synthesis between the placenta and embryo. Midgestation embryos are unable to engage the purine salvage pathway even when de novo purine synthesis is blocked either in vivo or in ex utero embryo culture, whereas placental tissue and trophoblasts retain the capacity to use either pathway. Disruption of de novo purine synthesis in mice causes reduced embryonic growth, impaired axial elongation, and abnormal brain and placental development, which are only partially rescued by supplementation with purine salvage precursors. In human placenta, trophoblast stem cells readily switch between the de novo and salvage pathways based on nutrient availability, and syncytiotrophoblasts (STB) preferentially rely on the salvage pathway. We identified guanosine monophosphate (GMP) as a metabolic checkpoint regulating STB differentiation, with insufficient GMP levels causing degradation of the small GTPase Rheb and failure of mTOR activation. Supplementation of purine salvage substrates restored GMP synthesis and STB differentiation in humans, but not mice. Further, in vivo measurements in humans revealed that maternal circulating hypoxanthine decreases during pregnancy and is further reduced in women with clinically small placentas, highlighting the role of hypoxanthine in supporting placental growth. These results uncover compartmentalized purine salvage between the embryo and placenta as a mechanism that limits competition for biosynthetic resources and enables coordinated growth during mammalian development.

## Introduction

Metabolism supports essential functions during mammalian development through mechanisms beyond energy production and biomass assimilation^5^. Metabolic gradients and spatiotemporal waves of metabolic activity are critical for development of the anterior-posterior axis formation^6,7^ and primitive streak elongation^8^, respectively. Moreover, metabolic activity plays a mechanistic role in morphogenesis and cell fate acquisition by controlling post-translational modifications^9^ and epigenetic regulation^10,11^. Although there are clear metabolic requirements (e.g. folate) for specific developmental processes, the degree to which individual metabolic pathways are expendable remains unclear.

As cells differentiate, their metabolic activities become constrained to support the functions of a particular lineage, suggesting that metabolic requirements are both cell type specific and temporally regulated during development. At the same time, cells retain a degree of metabolic plasticity that allows them to circumvent metabolic deficiencies, a capacity that is temporally regulated and varies within cell types depending both on gene expression and nutrient availability^5^. In mice, genetically induced metabolic deficiencies frequently result in embryonic maldevelopment or lethality, yet only rarely are placental phenotypes described^4,12,13^. In humans, inborn errors of metabolism are typically diagnosed in neonates after the placenta has been discarded, leaving an important gap in our understanding of how metabolic deficiencies influence placental development as well as the capacity for metabolic flexibility within placental and fetal compartments.

Midgestation murine development is a dynamic period marked by rapid growth of both the placenta and embryo, increased oxygen availability, and organogenesis^4^. With the essential function of transporting nutrients from maternal to fetal circulation, the placenta must balance its own growth while simultaneously supporting growth of the embryo. It remains unclear how the placenta achieves this rapid growth without directly competing with the embryo for biosynthetic resources. Our previous work identified coincident but distinct metabolic transitions in mouse placenta and embryo from gestational day (GD) 10.5-11.5, characterized by a sustained increase in purine synthesis in the embryo and a transient increase in arginine metabolism in the placenta^4^. Stable isotope tracing revealed rapid de novo purine synthesis in both the placenta and embryo, with embryonic tissues showing high de novo purine synthesis supported by maternally derived glucose and glutamine. The distinctions in purine synthesis between the placenta and embryo raise additional questions about how the placenta meets its nucleotide demands and its reliance on specific nutrients to its support growth and development.

Purine metabolites are critical for DNA and RNA synthesis, cell signaling pathways, and energy production, making this pathway of particular interest during midgestation, when the embryo and placenta are in a phase of exponential growth and morphogenesis. Cells produce purine monophosphates (e.g. Inosine monophosphate, IMP; Guanosine monophosphate, GMP; Adenosine monophosphate, AMP) through either a de novo synthesis or a salvage pathway. Tissue specific reliance on de novo versus salvage purine synthesis has been observed in adult mice^3^, raising questions about when such tissue specificity emerges during development and whether placental and embryonic tissues preferentially utilize distinct purine synthesis pathways. Here, we investigate purine synthesis metabolism during midgestation in placental and embryonic compartments, focusing on the capacity of these tissues to switch between de novo and salvage synthesis and its compatibility with developmental progression.

### Placental use of the purine salvage pathway

Our previous work used in vivo stable isotope tracing to demonstrate that midgestation embryos use maternally derived glucose and glutamine to rapidly synthesize purine monophosphates like IMP, GMP, and AMP (Figure 1A)^4^. Total enrichment of ^13^C from circulating glucose was equivalent between embryo and placenta in the purine precursor ribose 5-phosphate, but downstream enrichment of IMP was close to 60% in the embryo and only 40% in the placenta after a 4-hour infusion^4^. This data implied that although the placenta synthesizes the ribose backbone of purines using glucose from maternal circulation, it also uses alternative nutrient sources to generate the purine ring of IMP. This observation prompted us to investigate the use of the purine salvage pathway by the mouse placenta at midgestation (Figure 1A). Despite a sustained increase in purine metabolites in the embryo starting at GD10.5^4^, purine nucleosides and bases used by the salvage pathway like inosine, guanosine, guanine, and hypoxanthine are more abundant in placenta relative to the embryo from GD10.5 to GD13.5 (Figure 1B, S1A). Hypoxanthine, a metabolite at the intersection of purine salvage and catabolism that can be converted to IMP or broken down to xanthine and urate for excretion, is significantly more abundant in placenta compared to embryo throughout midgestation (Figure 1B, S1A) suggesting the placenta may use hypoxanthine to fuel the purine salvage pathway.

**Figure 1.**
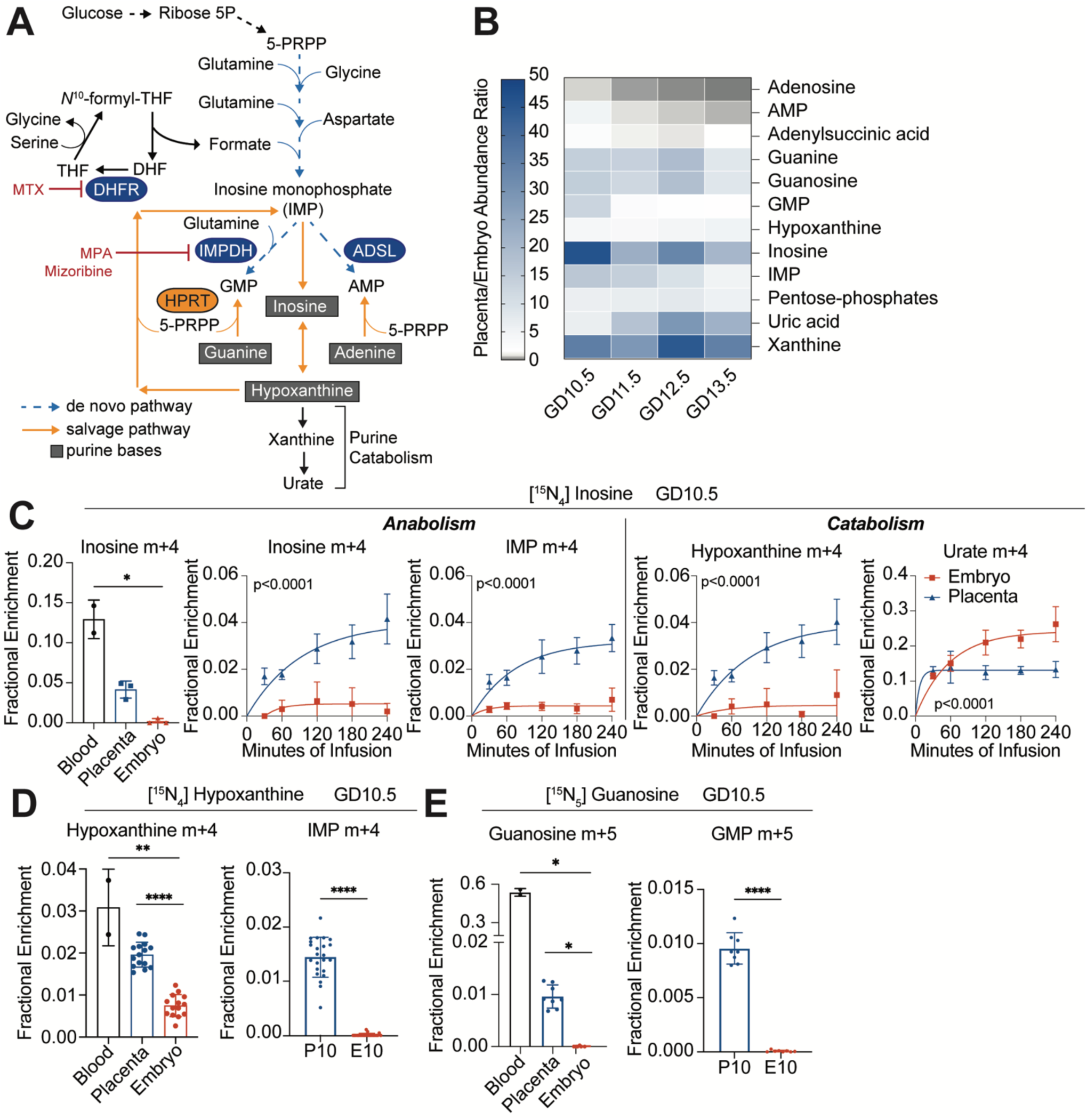
Assessment of salvage purine synthesis pathway in embryo and placenta. **A.** Schematic of de novo and salvage purine synthesis pathway. Methotrexate (MTX) inhibits purine de novo synthesis by targeting DHFR. Mycophenolic acid (MPA) or mizoribine (Miz) inhibits GMP synthesis by targeting IMPDH. **B.** Heatmap of purines, plotted as fold change placenta relative to paired embryo. **C**. [^15^N_4_] inosine enrichment in the blood, embryo and placenta at 180min during serial caesarian-section procedure (left). Fractional enrichment of inosine m+4, IMP m+4, hypoxanthine m+4 and urate m+4 are shown during infusion. **D**. [^15^N_4_] hypoxanthine enrichment in the blood, embryo and placenta (left). Fractional enrichment of IMP m+4 in placenta (P) and embryo (E) on GD10.5. **E**. [^15^N_5_] guanosine enrichment in the blood, embryo and placenta (left). Fractional enrichment of GMP m+5 in placenta (P) and embryo (E) on GD10.5. Statistical tests: (**C**) Kruskal-Wallis test followed by Dunn’s multiple-comparisons adjustment (Inosine m+4 enrichment in blood, embryo and placenta); plateau followed by one-phase association least-squares fitting. (**D**) Ordinary One-way ANOVA followed by Tukey’s multiple-comparisons adjustment (Hypoxanthine m+4 enrichment in blood, embryo and placenta); paired, two-tailed t test (IMP m+4). (**E**) Kruskal-Wallis test followed by Dunn’s multiple-comparisons adjustment (Guanosine m+5 enrichment in blood, embryo and placenta); Wilcoxon matched-pairs signed rank test(GMP m+5). ****: P < 0.0001; ***: P < 0.001; **: P < 0.01, *: P < 0.05, ns: P>0.05

To compare salvage pathway activity between placenta and embryo, we infused GD10.5 pregnant mice with [^15^N_4_]Inosine, [^15^N_4_]Hypoxanthine, or [^15^N_5_]Guanosine (Figure 1C-E). For inosine infusions, we performed serial cesarean-section procedures^4^ where a conceptus (placenta and embryo) was dissected every 30-60 minutes during a 4 hour infusion to assess the kinetics of transport and utilization. After 4 hours, the enrichment of circulating [^15^N_4_]Inosine was between 10-15% with 5% enrichment in placenta and close to no enrichment in embryo. The placenta showed accumulation of [^15^N_4_]Inosine throughout the procedure, which was both incorporated into IMP m+4 and converted to hypoxanthine m+4; whereas the embryo showed minimal accumulation of [^15^N_4_]Inosine or IMP m+4 but showed rapid synthesis of urate m+4 indicating labeled inosine was quickly catabolized in the embryo (Figure 1C). Infusion with [^15^N_4_]Hypoxanthine (Figure 1D, S1B) showed that enrichment of the tracer in placenta (2%) was close to enrichment values in circulation (3%) suggesting that the placenta efficiently takes up hypoxanthine from maternal circulation. Like inosine tracing, [^15^N_4_]Hypoxanthine taken up by placenta was used to synthesize IMP m+4 (1.5%) and catabolized to xanthine m+4 (2%) and urate (5%) (Figure S1B); whereas embryos showed no IMP m+4 but rapid production of xanthine m+4 (2.5%) and urate (10-15%) (Figure S1B). [^15^N_5_]Guanosine infusions had the highest level of circulating enrichment (50%) but showed only 1% enrichment in placental tissue suggesting that guanosine is not readily taken up by the placenta (Figure 1E). Despite this, the placenta effectively converted labeled guanosine into GMP m+5 (1%) with less catabolism to guanine (0.4%), and the embryo only catabolized [^15^N_5_]Guanosine to urate m+4 (15%) (Figure S1C). These results demonstrate the placenta uses circulating salvage precursors to generate purine monophosphates and this capacity is lacking in the embryo.

### Embryos cannot use the purine salvage pathway

Given that salvage precursor availability to the embryo is dependent upon placental transport, we tested the capacity for embryos grown in ex utero culture to use hypoxanthine to synthesize purines independent of placental influence. Embryos were dissected from pregnant mice on GD7.5 and grown for 3 days (comparable to GD10.5) as described previously^14^. Ex utero culture media (EUCM) was changed on culture day +3 to include [^15^N_2_]Glutamine with or without 20μM hypoxanthine. Excess hypoxanthine did not change glutamine uptake (Figure S2A) or increase embryonic hypoxanthine, IMP (Figure 2A), GMP, or AMP (Figure S2B) levels but did increase xanthine abundance (Figure 2A) suggesting no anabolic activity only robust hypoxanthine catabolism. Consistent with this, incubation of ex utero embryos with [^15^N_4_]Hypoxanthine showed only conversion to ^15^N urate (Figure S2C). Further, we observed no difference in the use of [^15^N_2_]Glutamine to support de novo synthesis when hypoxanthine was available (Figure 2B). These results support the conclusion that the embryo does not use the salvage pathway at this developmental stage. To test whether further development enables use of the salvage pathway by embryos, we performed [^15^N_4_]Hypoxanthine in vivo infusions on GD14.5, when placental maturation is complete. These data showed the placenta maintains hypoxanthine uptake throughout gestation, but uptake decreases over time in the embryo, which corresponds with reduced enrichment in xanthine m+4 and urate m+4 in the embryo at GD14.5 (Figure S2D). Collectively this data indicates that during a phase of rapid growth, there are compartment specific strategies to synthesize purines where the embryo relies only on de novo purine synthesis and tightly controls hypoxanthine levels by catabolism to xanthine and urate.

**Figure 2.**
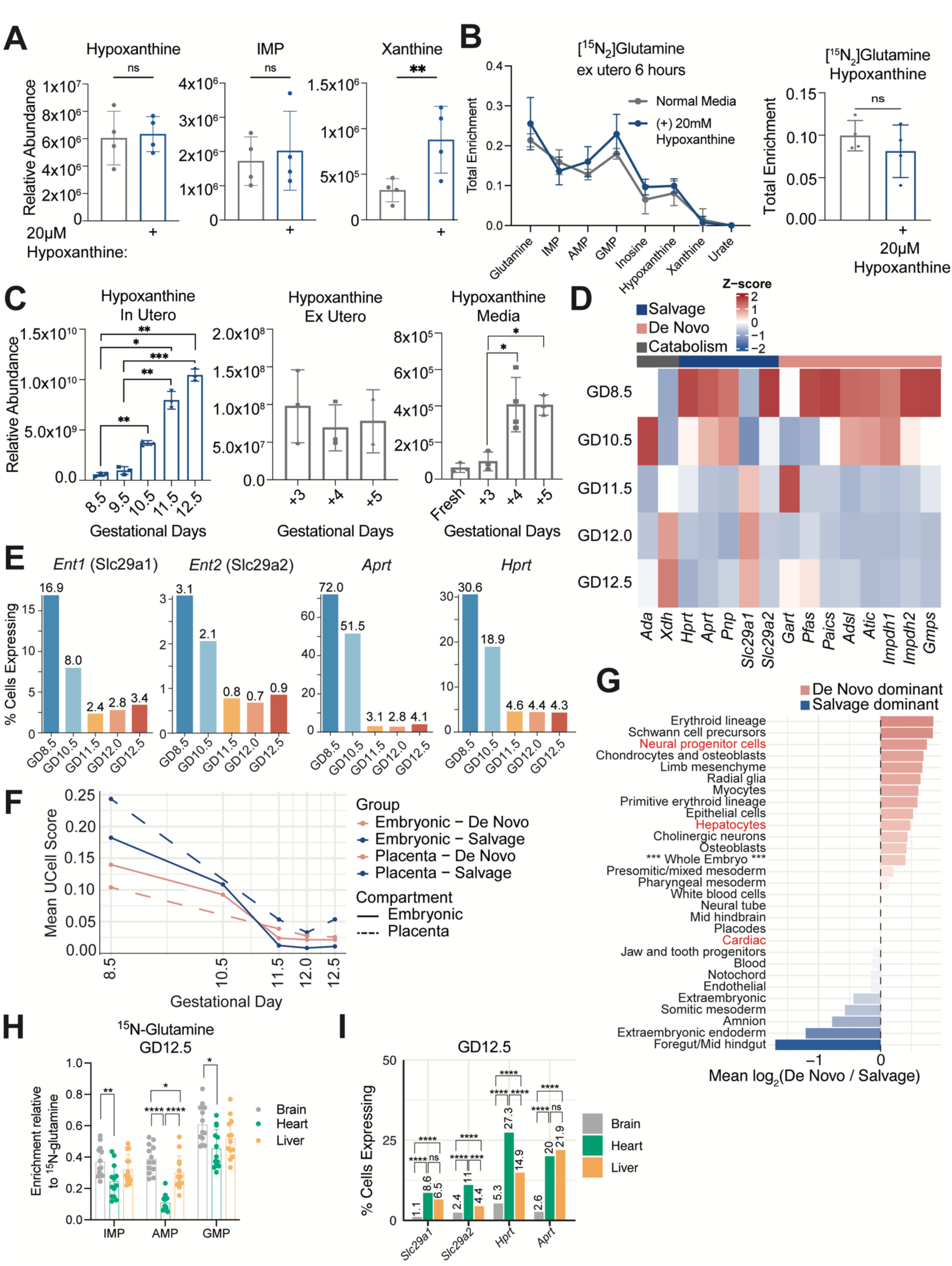
De novo purine synthesis pathway is dominant in embryo. **A**. Relative abundance of hypoxanthine, IMP and xanthine in ex utero embryo treated with or without 20μM hypoxanthine for 6h on GD10.5. **B**. Total enrichment of indicated metabolites from 6h-[^15^N_2_]Glutamine tracing in E10.5 ex utero embryo treated with or without 20μM hypoxanthine (left). Total enrichment of hypoxanthine with or without 20 µM hypoxanthine supplementation (right). **C**. Hypoxanthine abundance from E8.5 to E12.5 in utero embryos (left) and E10.5 to E12.5 ex utero embryo from (middle). Hypoxanthine abundance in conditioned media of ex utero embryo from E10.5 to E12.5(right). **D**. Heatmap of Z-scored mean expression of purine metabolism genes from E8.5–E12.5 in utero whole embryo scRNAseq data. Z-scores were computed per gene across stages. n = 226,257 total cells (4,349 at E8.5; 21,546 at E10.5; 39,313 at E11.5; 80,128 at E12.0; 80,921 at E12.5). **E** Percentage of cells expressing key purine salvage and transport genes across E8.5–E12.5 from whole embryo in utero scRNAseq data. Numbers above bars indicate the percentage of cells with detectable expression (normalized expression > 0). **F**. UCell pathway activity scores for salvage and de novo purine synthesis pathways in embryonic versus extraembryonic/placental tissues from E8.5–E12.5. Lines represent mean UCell scores per compartment; solid lines indicate embryonic tissues and dashed lines indicate extraembryonic tissues. Gene signatures: salvage (Hprt, Aprt, Pnp) and de novo (Gart, Pfas, Paics, Adsl, Atic). Extraembryonic tissues maintain salvage pathway dominance (salvage > de novo) at every developmental stage (Wilcoxon signed-rank test, P < 0.05 at all stages). Embryonic tissues switch from salvage-dominant at E8.5 to de novo-dominant after E10.5 (P < 2.2 × 10⁻¹⁶ at E11.5–E12.5), cell counts n = 222,489 embryonic and 3,768 extraembryonic cells. **G**. Mean log₂(de novo / salvage) expression ratio for each tissue type across all developmental stages in the in utero scRNAseq dataset. Mean expression of salvage pathway genes (Hprt, Aprt, Pnp) and de novo pathway genes (Gart, Pfas, Paics, Adsl, Atic) was computed per tissue per stage, and the log₂ ratio was averaged across stages. Negative values (blue) indicate salvage pathway dominance; positive values (red) indicate de novo dominance. Tissues required ≥100 cells per stage to be included; minimum 50 cells per tissue-stage combination for ratio calculation. **H**. Total Enrichment of IMP, AMP and GMP relative to [γ-^15^N]Glutamine in tissue from in vivo infusion. **I**. Expression of purine salvage and nucleoside transport genes from scRNAseq mouse embryo atlas^29^ at E12.5. Bar plots show the percentage of cells expressing Slc29a1 (Ent1), Slc29a2 (Ent2), Hprt and Aprt in Brain (CNS neurons, n=35,978 cells), Heart (Cardiomyocytes, n=245 cells), and Liver (Hepatocytes, n=1,492 cells). Pairwise comparisons were performed using the Wilcoxon rank-sum test (two-sided) on per-cell expression values, with Benjamini-Hochberg correction for multiple testing (12 tests). Statistical tests: (**A**) Unpaired, two-tailed t test; (**B**) Two-way ANOVA with Sidak’s multiple comparisons (left) and unpaired, two-tailed t test (right); (**C-H**) Ordinary One-way ANOVA with Tukey’s multiple comparisons; (I) Wilcoxon rank-sum test (Mann-Whitney U). ****: P < 0.0001; ***: P < 0.001; **: P < 0.01, *: P < 0.05, ns: P>0.05

Interestingly, even when cultured in excess hypoxanthine, midgestation embryos displayed 10% ^15^N enrichment in hypoxanthine during [^15^N_2_]Glutamine tracing (Figure 2B). Metabolomics of in utero embryos from GD8.5 to GD12.5 shows a significant increase in hypoxanthine levels during the midgestation metabolic transition (Figure 2C). Ex utero embryos do not show an increase in hypoxanthine from culture days +3 to +5 (comparable to GD10.5-GD12.5) but excretion of hypoxanthine in the media increases significantly between culture days +4 and +5 (Figure 2C). This data suggests that the embryo tightly maintains hypoxanthine levels and excretes hypoxanthine into fetal circulation where it can be taken up by the placenta and used to support purine salvage, converted to urate, or excreted into maternal circulation.

To better understand mechanisms by which purine synthesis is regulated in the embryo we performed single cell RNA sequencing (scRNAseq) on in utero embryos at GD8.5 and GD10.5-12.5 to observe changes in metabolic gene expression. Comparing the expression of enzymes in de novo and salvage purine synthesis as well as purine catabolism showed a rapid decline in salvage enzyme expression during the midgestation metabolic transition (Figure 2D). Specifically, the number of cells expressing Equilibrative nucleoside transporters 1 and 2 (*Ent1/Ent2*) encoding Slc29a1/2 and the salvage enzymes Adenine Phosphoribosyltransferase (*Aprt*) and Hypoxanthine/Guanine Phosphoribosyltransferase (*Hprt*) significantly declined between GD10.5 and GD11.5 (Figure 2E). The overall mean UCell score of expression of purine de novo and salvage enzymes shifted in the embryo between GD10.5 and GD11.5 from higher salvage at GD8.5 to higher de novo at GD12.5, while extra embryonic tissues (GD8.5) and the placenta (GD10.5-GD12.5) maintained high salvage expression throughout midgestation (Figure 2F). This data is consistent with our infusion data showing more robust purine salvage occurring in placental tissues compared with embryonic tissues.

Given that this transition occurs at the beginning stage of organogenesis, we compared the gene expression profiles of different embryonic and extraembryonic tissues across developmental stages GD8.5-GD12.5 to define cell types that predominantly use the de novo pathway or the salvage pathway. This data shows gastrointestinal cells and somitic mesoderm as the only embryonic cell types that predominately express purine salvage enzymes with other cell types expressing enzymes in either both pathways or primarily de novo enzymes (Figure 2G). We infused mice at GD12.5 with [γ-^15^N]Glutamine to determine the degree of de novo purine synthesis in fetal organs. Comparison of ^15^N enrichment in IMP, AMP, and GMP in fetal brain, liver, and heart showed the highest ^15^N enrichment in brain and the lowest ^15^N enrichment in heart (Figure 2H). This data is consistent with the gene expression profiles indicating predominant use of the de novo pathway (Figure 2G) in fetal brain and use of both pathways in fetal heart, which displays higher expression of purine salvage pathway genes (Hprt, Aprt) and nucleoside transporter genes (Slc29a1/Ent1, Slc29a2/Ent2) at gestational day 12.5 than brain or liver (Figure 2I). Collectively, this data indicates that cell-type specific strategies for purine synthesis are defined early during development driven by gene expression changes across the whole embryo and constraint of purine metabolism during the midgestation metabolic transition (GD10.5-11.5).

Our data indicate that under normal developmental conditions, the midgestation embryo shows minimal use of the salvage pathway, so we tested the capacity to engage purine salvage when the de novo synthesis pathway was blocked. We first treated GD10.5 pregnant dams with the dihydrofolate reductase (DHFR) inhibitor (Figure 1A), methotrexate (10mg/kg) one hour prior to [^15^N_4_]Hypoxanthine infusion. Methotrexate (MTX) treatment increased hypoxanthine m+4 and IMP m+4 enrichment in the placenta however the embryo showed reduced hypoxanthine uptake and no enrichment in IMP (Figure 3A). This suggests that with acute blockade of de novo purine synthesis, the embryo does not compensate with increased purine salvage. To determine if longer term treatment with MTX enables embryonic compensation, we treated GD10.5 mice with once daily MTX (5mg/kg) and then collected tissues on GD12.5 to assess purine abundance. After 2-day treatment, we saw decreased levels of IMP, GMP, and AMP in the embryo (Figure 3B) whereas only AMP levels were altered in placental tissue (Figure S3A). This demonstrates the capacity for placental tissue to adapt to deficiency in de novo purine synthesis by compensating with the salvage pathway, while the embryo lacks this metabolic flexibility. This suggests that the constraint on purine salvage driven by reduced enzyme expression limits the embryo’s capacity to engage compensatory pathways and circumvent metabolic deficiencies.

**Figure 3.**
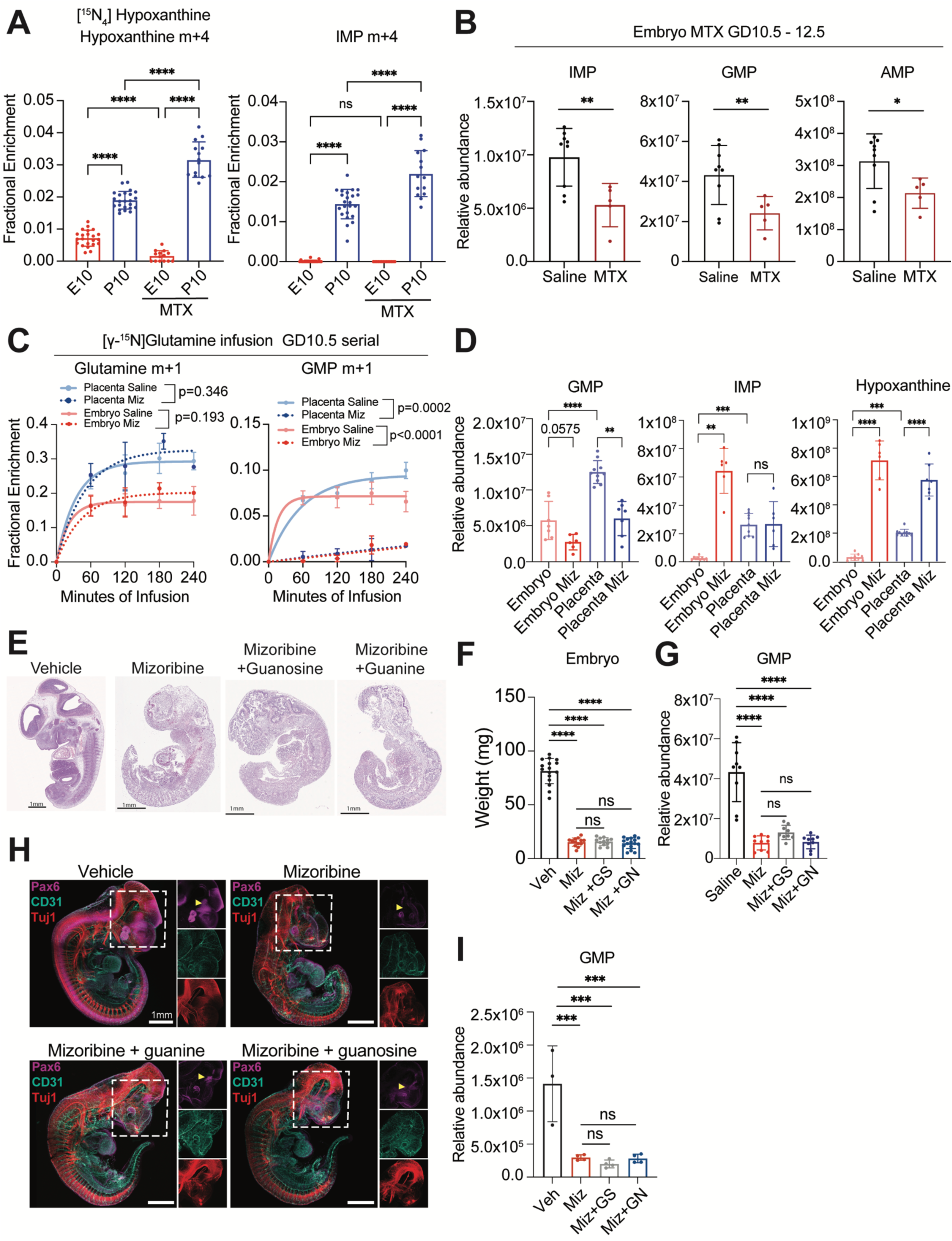
Embryos are unable to utilizing purine bases for salvage synthesis. **A**. 10mg/kg methotrexate (MTX) pretreatment (1h) with [^15^N_4_] hypoxanthine infusion. Fractional enrichment of hypoxanthine m+4 and IMP m+4 in embryo and placenta. **B**. Relative abundance of IMP, GMP and AMP in GD12.5 embryo from mice treated with 5mg/kg MTX from GD10.5-GD12.5. **C**. Mizoribine(50mg/kg) treatment (1h) prior to [γ-^15^N] Glutamine serial caesarian-section infusion (left) and GMP m+1 (right) in embryo and placenta at indicated time points MIZ=mizoribine. **D**. Relative abundance of GMP, IMP and hypoxanthine at 180min during serial caesarian-section procedure. **E**. Representative HE staining showed morphology of embryos at E12.5 with indicated treatment from E10.5 to E12.5. Mizoribine: 50mg/kg. Guanosine: 8mg/kg. Guanine: 5mg/kg. **F**. Embryo weights after 2 day treatment. GS: Guanosine. GN: Guanine. **G**. Relative abundance of GMP in embryo after 2 day treatment. **H**. Whole-embryo immunostaining for PAX6, βIII-tubulin (Tuj1), and CD31 in ex utero grown embryos treated with 50μM Mizoribine with and without 50μM guanosine or guanine. **I**. Relative abundance of GMP in ex utero embryos with indicated treatment. Statistical tests: One-way ANOVA tests, Dunnett’s multiple comparison test (**A**, **D, G, I)**, **(F)** Welch’s one way, Dunnett’s T3 multiple comparison. (**C**) Plateau followed by one-phase association least-squares fitting, Extra sum-of-squares F test; Unpaired, two-tailed t test (**B**). ****: P < 0.0001; ***: P < 0.001; **: P < 0.01, *: P < 0.05, ns: P>0.05

Since altered AMP and adenosine metabolism can influence cellular energy balance^15^ and vascular biology^16,17^ in addition to reducing precursors for DNA and RNA synthesis, we tested whether just blocking de novo GMP synthesis impairs embryonic development and if supplementation with salvage metabolites guanosine or guanine could maintain GMP levels. We treated GD10.5 pregnant mice with mizoribine, an inhibitor to inosine monophosphate dehydrogenase (IMPDH) to block de novo synthesis of GMP (Figure 1A) one hour prior to serial caesarean infusion with [γ-^15^N]Glutamine. Our previous data^4^ demonstrated reduced labeling in glutamine in embryo relative to placenta even at steady state conditions. Treatment with mizoribine did not alter glutamine uptake rates in either tissue but did blunt ^15^N enrichment in GMP (Figure 3C) consistent with the drug working as expected. Total abundance of GMP was significantly decreased in both embryo and placenta with mizoribine treatment (Figure 3D) indicating that endogenous guanine and guanosine salvage is not sufficient to maintain GMP levels with acute IMPDH deficiency in the placenta or embryo. In embryos, IMP and hypoxanthine levels were significantly increased (Figure 3D) consistent with a compensatory increase in de novo IMP synthesis and breakdown to hypoxanthine and xanthine compared to vehicle treated embryos (Figure S3B). The placenta displayed no overall increase in IMP levels but did show increased rates of de novo IMP synthesis relative to vehicle treated placenta, suggesting reduced GMP abundance increases de novo activity in both compartments (Figure S3B). Overall hypoxanthine levels were significantly increased in placenta relative to vehicle treated placenta (Figure 3D) and displayed increased ^15^N hypoxanthine m+1 and ^15^N xanthine m+1 synthesis (Figure S3B) suggesting either local IMP and inosine catabolism, or uptake of embryo-derived intermediates. These experiments demonstrate that the acute metabolic response to IMPDH blockade in developing tissues is compensatory de novo activity as well as purine cycling.

We next tested the capacity for either compartment to increase purine salvage using supplemented guanosine or guanine to overcome longer IMPDH deficiency and continue developmental progression. We treated GD10.5 pregnant mice for 2 days with 50mg/kg mizoribine with and without supplementation of 8mg/kg guanosine or 5mg/kg guanine. Histological examination of embryos treated with mizoribine displayed pronounced developmental abnormalities including markedly reduced size (Figure 3E), disorganized tissue architecture, impaired axial elongation, and poorly defined organ boundaries. Brain development was greatly impaired, particularly at the forebrain region, somite organization was compromised, and limb bud outgrowth was markedly reduced, consistent with a broad impairment of growth and tissue patterning during organogenesis. Embryos co-treated with mizoribine and guanine or guanosine showed no restoration of embryo size or morphology (Figure 3F), and consistent with our in vivo [^15^N_5_]Guanosine infusion data, supplementation with excess guanosine and guanine did not rescue GMP levels in the embryo (Figure 3G) with guanosine or guanine supplementation only resulting in a marked increase in guanine levels (Figure S3C). We similarly treated ex utero embryos with 50μM mizoribine with and without supplementation of 50μM guanine or 50μM guanosine which showed similar morphological phenotypes to in vivo embryos, along with reduced presence of red blood cells in the yolk sac vasculature upon mizoribine treatment, which was partially rescued by guanine supplementation (Figure S3E). Supplemented ex utero embryos exhibited evident abnormalities relative to controls, indicating that guanine supplementation only partially alleviates the effects of IMPDH inhibition. Whole-embryo immunostaining for neural progenitors, differentiated neurons, and vasculature revealed that Mizoribine significantly reduced the Pax6⁺ progenitor pool in the brain and spinal cord, whereas Tuj1⁺ neurons and CD31⁺ endothelial cells were largely preserved. Although supplementation with guanine or guanosine did not fully restore the Pax6⁺ progenitor population, a slight expansion of the Pax6⁺ domain in the embryonic forebrain was observed (Figure 3H). As with our in vivo data, supplementation of ex utero embryos with guanine and guanosine did not rescue GMP levels (Figure 3I, S3D). These data further demonstrate de novo GMP synthesis is required for proper overall embryonic growth and specifically development of forebrain region and red blood cells. Additionally, it suggests that when de novo GMP synthesis is blocked, most cell types cannot engage purine salvage pathways even when bases are provided in excess.

### Syncytiotrophoblast differentiation constrains purine synthesis

Mizoribine treatment for 2 days also influenced placental growth and development. Histological examination of placentas treated with mizoribine indicated a reduction in the labyrinth (L) region relative to junctional zone (JZ) (Figure 4A-B). Overall tissue weight was reduced with neither phenotype rescued by excess guanine or guanosine (Figure 4C). Similar to MTX treatment, placentas treated with mizoribine for 2 days showed metabolic compensation and were able to maintain GMP levels even without guanine or guanosine supplementation (Figure 4D). Despite maintenance of GMP abundance, mizoribine-treated placentas showed alterations in markers of specific cell populations that likely explains the altered L/JZ ratio. The most striking finding was a significant reduction in *Gcm1* and *Synb,* markers of the syncytiotrophoblast II (SynII) layer that directly contacts fetal endothelial cells and facilitates nutrient transfer from maternal blood to fetal circulation^18^ (Figure 4E). This reduced expression could not be rescued by guanine or guanosine supplementation and presumably contributes to the reduced placental size. The reduction in SynII cells were accompanied by an increase in *Tpbpa* and *Ctsq* expression, markers of spongiotrophoblast cells that make up the junctional zone (Figure 4E). Unlike the reduction in SynII genes, increased expression of spongiotrophoblast genes were rescued to vehicle levels upon guanine supplementation. These data suggest that purine metabolism has a functional role in cell lineage determination in the placenta with mechanisms that may vary by cell type. Interestingly, guanine supplementation seemed to increase stem cell and syncytiotrophoblast I markers in the presence of mizoribine (Figure 4E), which raises additional questions about how purine availability influences placental development.

**Figure 4.**
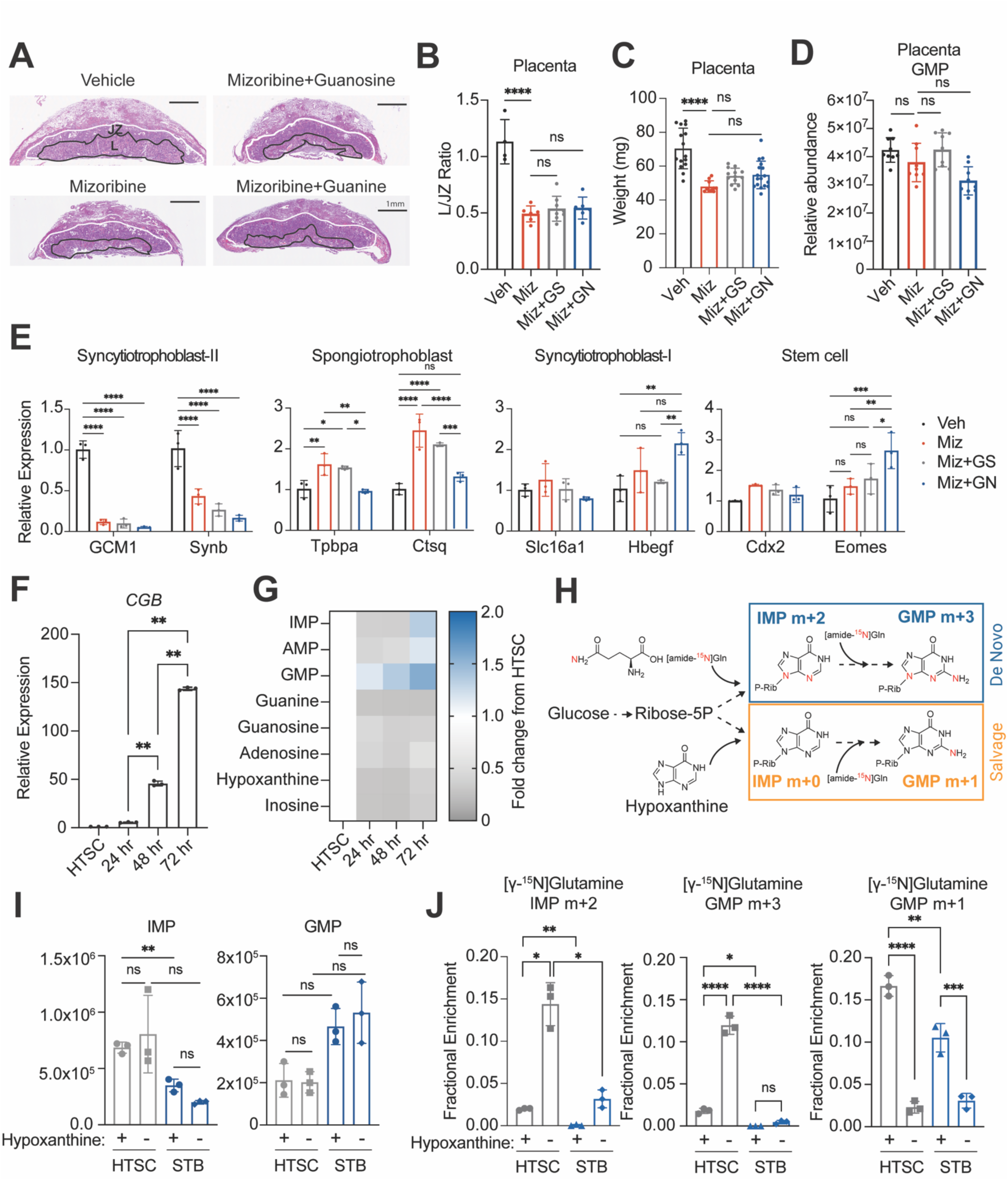
Placental tissue can utilize purine bases for salvage synthesis. **A**. Representative HE staining showed morphology of placentas at E12.5 with indicated treatment from GD10.5 to GD12.5. Mizoribine: 50mg/kg. Guanosine:8mg/kg. Guanine: 5mg/kg. Labyrinth layer (L) was marked with black line and junctional zone (JZ) was area between white line and black line. **B**. Quantification of labyrinth layers and junctional zone area. Area measurement was done by ImageJ/Fiji software. **C**. Placenta weight at GD12.5 with indicated treatment from GD10.5 to GD12.5. **D**. Relative abundance of GMP in GD12.5 placenta with indicated treatment from GD10.5 to GD12.5. **E**. RT-qPCR analysis of cell type-specific maker genes in GD12.5 placenta with indicated treatment from GD10.5 to GD12.5. **F**. RT-qPCR analysis of human syncytiotrophoblast (STB) maker *CGB* at indicated time points during human trophoblast stem cell (HTSC) differentiation. **G**. Heatmap of purine metabolites at indicated time points during HTSC differentiation. **H**. Schematics illustrating ^15^N labeling of GMP from [γ-^15^N] Glutamine in HTSC and STB cultured in media with or without hypoxanthine. **I**. Relative abundance of IMP or GMP in HTSC and STB cultured in media with or without hypoxanthine. STB differentiation was performed in media without hypoxanthine, which was added for the last 6 hours. **J**. Fractional enrichment of IMP m+2, GMP m+3 and GMP m+1from [γ-^15^N] Glutamine in HTSC and STB cultured in media with or without hypoxanthine. Statistical tests: (**B-D)** One-way ANOVA tests, Tukey’s multiple comparisons; (**E**) Two-way ANOVA tests; (**F, I**) Welch’s one-way ANOVA, Dunnett’s T3 multiple comparisons tests; (**J**) Welch’s one way ANOVA, Dunnetts tests for IMP m+2, One-Way ANOVA, Tukey’s multiple comparisons tests for GMP m+3 and GMP m+1. ****: P < 0.0001; ***: P < 0.001; **: P < 0.01, *: P < 0.05, ns: P>0.05

To better understand the dynamics of purine metabolism and trophoblast cell fate, we turned to cultured human trophoblast stem cells that can be differentiated to syncytiotrophoblasts (STBs) with the addition of forskolin, which leads to cellular fusion to form a multinucleated syncytium that produces the pregnancy hormone chorionic gonadotropin beta (*CGB*)^19^. Induction of *CGB* and other STB marker genes, Syndecan 1 (*SDC1*), and Aromatase (*Cyp19A1)* were observed within 24-48 hours after forskolin treatment of HTSCs (Figure 4F, S4A). Metabolomics analysis during STB differentiation showed IMP and AMP levels acutely drop and then by 72 hours rise to levels above HTSCs (Figure 4G). Interestingly, GMP levels steadily increase each day during HTSC differentiation to STBs (Figure 4G). To test the preferential use of the de novo versus purine salvage pathways in each cell type, we incubated HTSCs or differentiated STBs (5-day differentiation) with [γ-^15^N]Glutamine. Since the culture media for these cells contains hypoxanthine, we performed these tracing experiments with and without hypoxanthine in the media to assess if nutrient availability altered the use of either pathway. If cells use de novo synthesis, [γ-^15^N]Glutamine will generate IMP m+2 and GMP m+3 whereas if cells use unlabeled hypoxanthine to generate IMP via purine salvage, then GMP m+1 will be observed (Figure 4H). In the absence of hypoxanthine both HTSCs and STBs maintained levels of IMP and GMP indicating that hypoxanthine availability is not necessary for HTSC or STB maintenance of purine monophosphate pools (Figure 4I). Like what we observed in both in vivo and ex utero grown mouse embryos, hypoxanthine levels seem to be tightly controlled in HTSCs and in STBs since hypoxanthine depletion in the media led to a significant increase in synthesis of ^15^N hypoxanthine (m+2) but no overall change in hypoxanthine levels in either HTSCs or STBs (Figure S4B). Removal of hypoxanthine from the media did not increase glutamine uptake in either cell type (Figure S4C) but did result in a significant increase in IMP m+2 and GMP m+3 enrichment and decrease in GMP +1 enrichment in HTSCs indicating predominant use of de novo purine synthesis when salvage bases are not available (Figure 4J). When hypoxanthine was present in the media, HTSCs displayed reduced IMP m+2 and GMP m+3 and increased GMP m+1 indicating use of the purine salvage pathway (Figure 4J). This data demonstrates that HTSCs switch between use of the de novo and salvage synthesis pathways based on salvage precursor availability. Alternatively, STBs showed no enrichment in IMP m+2 or GMP m+3 but enrichment of GMP m+1when hypoxanthine was available indicating predominant use of the purine salvage pathway (Figure 4J). When hypoxanthine was removed, STBs showed a reduction in GMP m+1 and only a modest increase in IMP m+2 and GMP m+3 (Figure 4J). This data indicates that STBs are constrained to use the salvage pathway and have limited capacity to induce de novo synthesis when salvage precursors are not available. The lack of any change in GMP levels in the absence of hypoxanthine in STBs is interesting given there is only limited increase in de novo synthesis however this could be explained by reduction in GMP catabolism or conversion of GDP to GMP.

### GMP synthesis is necessary for trophoblast syncytialization

As HTSCs differentiate to STBs, GMP levels increase (Figure 4G), so we tested the impact of blocking de novo GMP synthesis in HTSCs and fully differentiated STBs using the IMPDH inhibitor, Mycophenolic Acid (MPA). Like mizoribine, MPA treatment blocks GMP synthesis from the de novo pathway and purine salvage by hypoxanthine (Figure 1A) since hypoxanthine is first converted to IMP via the salvage enzyme Hypoxanthine phosphoribosyl transferase 1 (HPRT1). Treatment of HTSCs and STBs with MPA in complete media (containing hypoxanthine) reduced overall GMP levels consistent with the drug working as expected (Figure 5A). Tracing with [γ-^15^N]Glutamine showed reduced GMP m+1, and tracing with [^15^N_4_]hypoxanthine showed reduced labeling in GMP m+4 as expected for inhibition of IMPDH (Figure 5A, S5B). Compensatory increase in IMP pool size was only seen in HTSC, whereas IMP m+2 indicating de novo synthesis was increased in both HTSCs and STBs (Figure S5A). This is consistent with the compensatory response we observed in mizoribine treated embryos and placenta (Figure 3D, S3B).

**Figure 5.**
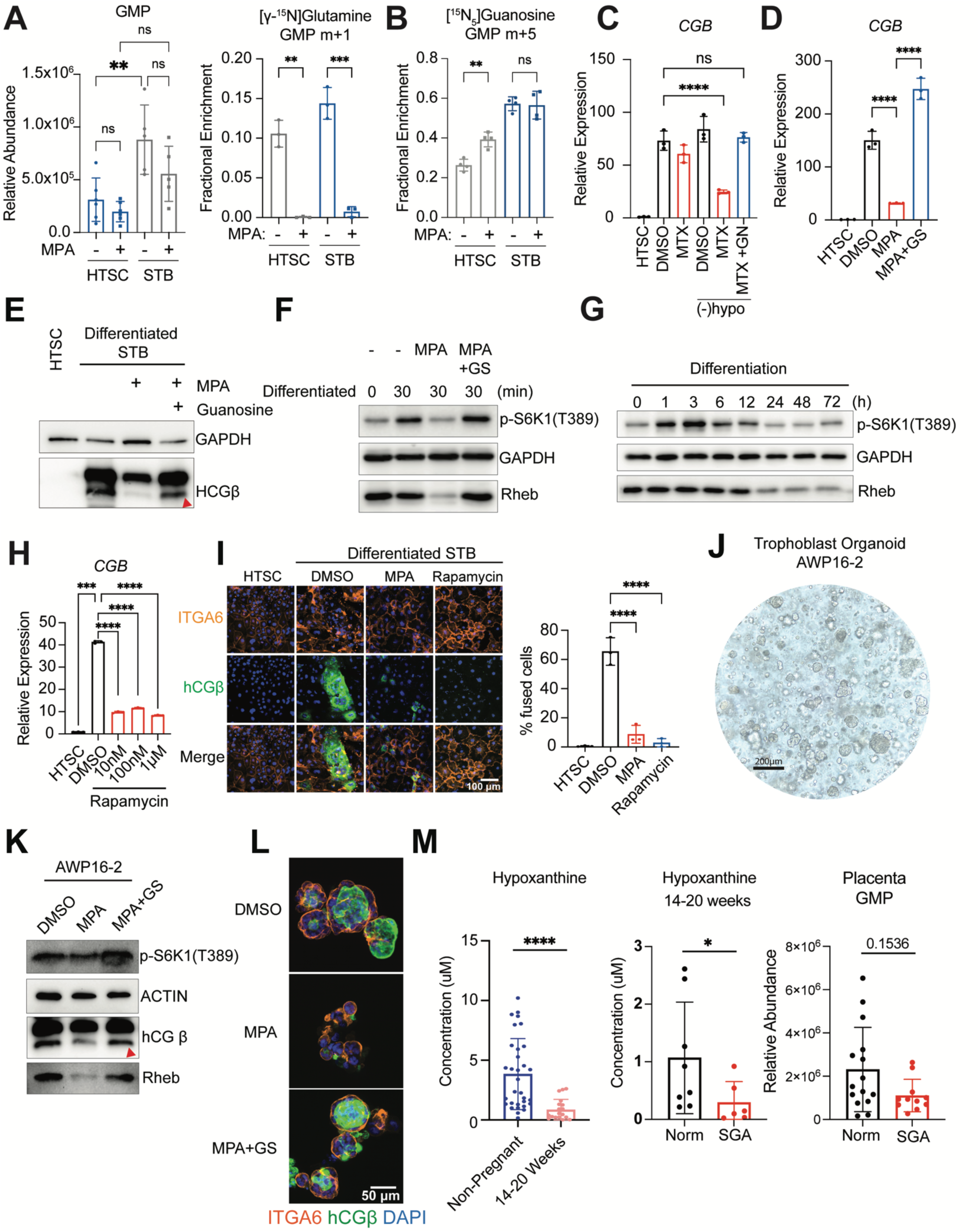
GMP synthesis is essential for placental development. **A**. Relative abundance of GMP in HTSC and STB after treating with 5 μM MPA for 24h (left). Fractional enrichment of GMP m+1 from [γ-^15^N] Glutamine after 24h-MPA treatment (right). **B**. Fractional enrichment of GMP m+5 from [^15^N_5_] Guanosine after 24h-MPA treatment. **C**. RT-qPCR analysis of *CGB* with indicated treatments. Treatments lasted for 5 days after differentiation initiated. MTX (10μM): methotrexate. Hypo: hypoxanthine. GN (20μM): guanine. **D**. RT-qPCR analysis of *CGB* with indicated treatments. Treatments lasted for 5 days after differentiation initiated. GS (20μM): guanosine. **E**. Immunoblots of hCGβ in differentiated HTSC with indicated treatments. Treatments lasted for 5 days after differentiation initiated. **F**. Immunoblots of p-S6K1(T389) and Rheb after differentiation for 30min in hTSC with indicated 24h pretreatments. **G**. Immunoblots of p-S6K1(T389) and Rheb at different time points during hTSC differentiation. **H**. RT-qPCR analysis of *CGB* with Rapamycin treatment, and treatments lasted for 5 days after differentiation initiated. **I**. Immunofluorescence staining of hCGβ (green) and ITGA6(Cytotrophoblast marker, orange) and cell fusion percentage calculation (# cells with multiple nuclei) of HTSC after 5 days of differentiation with indicated treatments. **J**. Brightfield image of term human trophoblast organoid AWP16-2. **K**. Immunoblots of hCGβ, p-S6K1(T389) and Rheb in AWP16-2 with indicated treatments for 72h. MPA: 5μM. Guanosine: 20 μM. **L**. Immunofluorescence staining of hCGβ (green) and ITGA6(orange) in organoid with indicated treatments for 96h. M. (left) Circulating hypoxanthine concentration in plasma from non-pregnant and pregnant patients(14-20 weeks); (middle) Circulating hypoxanthine level in pregnant patients (14-20 weeks) with normal or small for gestational age (SGA) placentas; Placental GMP level in patients with normal or SGA placentas. Statistical tests: **(A-D, H-I)** Ordinary One-way ANOVA, Tukey’s multiple comparisons tests; **(M)** Unpaired, two tailed t test (left), Unpaired, two tailed Welch’s t test (middle and right).

To interrogate the purine metabolism requirements during STB differentiation, we first tested whether limiting HTSCs to either the de novo or salvage pathways had an impact on STB differentiation. Removal of hypoxanthine from the media requires use of only the de novo synthesis pathway whereas treatment with methotrexate (MTX) forced only use of the purine salvage pathway (Figure 1A) with successful STB differentiation measured by induction of *CGB* (Figure 5C). Only when hypoxanthine was removed in the presence of MTX was *CGB* induction blocked suggesting that HTSCs reliant on only de novo purine synthesis or only purine salvage are equally competent to differentiate to STBs. We then tested the requirement for GMP synthesis in HTSCs treated with MPA when forskolin was added to induce STB differentiation and then assessed *CGB* induction after 5 days. MPA treatment blunted STB differentiation and this could be rescued completely by supplementation with guanosine at the time of MPA treatment (Figure 5D-E). Although MPA reduced HTSC proliferation, IMPDH inhibition had no impact of viability of differentiated STBs, which excludes the possible influence from cell viability on STB differentiation (Figure S5D). This data indicates that GMP levels act as a metabolic checkpoint for human STB differentiation and that unlike mouse SynII differentiation, IMPDH inhibition in human trophoblasts can be overcome by supplementation with guanosine.

Shortly after implantation, trophoblasts begin to produce human chorionic gonadotropin (hCG), which that stimulates cytotrophoblasts to differentiate to STBs via the luteinizing hormone (LH)/hCG receptor, a G protein coupled receptor that stimulates cyclic AMP (cAMP) production^20^. Activation of this pathway is mimicked in cultured trophoblasts by forskolin treatment and can activate PKA-dependent and -independent pathways, the latter of which can activate the MAPK/ERK pathway^20^. We tested whether MPA blocks activation of ERK acutely after forskolin treatment and observed no difference in phosphorylated ERK at 10 and 30 minutes between MPA pretreated and vehicle treated cells suggesting that MPA blocks STB differentiation downstream of ERK. (Figure S5C). ERK activation can lead to downstream activation of mTORC1^21^, and purine deprivation has been shown to inhibit mTORC1^22,23^, so we measured phosphorylation of S6 Kinase (ph-S6K T389) as a marker of mTORC1 activity. We observed decreased ph-S6K in MPA pretreated HTSCs 30 minutes after forskolin addition that was rescued by supplementation with guanosine (Figure 5F). Decreased GMP levels have been shown to inhibit mTORC1 through degradation of the upstream GTPase, Rheb^22,23^ and we also observed decreased Rheb protein abundance in MPA treated HTSCs, which was also rescued by guanosine supplementation (Figure 5F). The role of mTORC1 in human STB differentiation has previously been studied in choriocarcinoma or immortalized trophoblast lines, which showed rapamycin treatment promoted syncytialization^24^. These studies showed that amino acid deprivation led to activation of AMP activated protein kinase (AMPK), and subsequent mTORC1 inhibition promoted macropinocytosis and trophoblast syncytialization 12 hours after forskolin treatment. To better understand the dynamics of mTORC1 activation during STB differentiation, we profiled ph-S6K and Rheb levels from 1 to 72 hours after forskolin treatment in first trimester primary human trophoblast stem cells^19^ and observed an acute activation of mTORC1 during the first 12 hours followed by decreased expression of Rheb and reduced ph-S6K from 24 to 72 hours (Figure 5G). This data suggested that mTORC1 activity is required for HTSC commitment to STB. To further test this, we treated HTSCs with increasing doses of rapamycin upon forskolin treatment to initiate STB differentiation. Rapamycin treatment blocked induction of *CGB* at both the mRNA and protein levels and prevented cellular fusion, phenocopying MPA treatment (Figure 5H-I). Collectively, this data demonstrates that GMP levels act as a checkpoint on initial mTORC1 activation during HTSC commitment to the STB lineage.

Finally, to test the universality of this mechanism in a mixture of cell types, we derived human placental organoids (AWP16-2) from healthy, term patients who did not experience labor or rupture of membranes and delivered by cesarean section (Figure 5J) and treated them with MPA with and without guanosine supplementation for 3 days. Treatment with MPA reduced Rheb protein levels, modestly reduced ph-S6K, and reduced hCGβ protein expression, all of which were rescued by exogenous guanosine supplementation (Figure 5K-L). Collectively, our data suggests that trophoblast stem cells have a GMP-mTORC1 metabolic checkpoint that ensures adequate purine availability prior to commitment to the STB lineage. Upon differentiation to STBs, cells are constrained to use hypoxanthine and the purine salvage pathway to maintain GMP levels. Since our in vivo and in vitro data highlight the importance of hypoxanthine as a metabolite that supports placental growth and development, we measured circulating hypoxanthine during early human pregnancy and found a significant decrease in pregnant individuals between 14-20 weeks relative to non-pregnant patients (Figure 5M, left). Interestingly, individuals who displayed small for gestational age (SGA) placentas where the placental weight was less than the 10^th^ percentile^25^ showed significantly lower levels of circulating hypoxanthine between 14-20 weeks relative to patients with normal placental size (Figure 5M, middle), and SGA placentas showed a modest reduction in placental GMP levels upon delivery (Figure 5M, right). This data suggests that insufficient circulating hypoxanthine levels early in pregnancy may limit GMP synthesis during human placental development leading to reduced placental growth throughout pregnancy.

## Discussion

Metabolic functions support many aspects of mammalian development. Genetic, pharmacologic, or nutritionally induced defects in metabolic pathways result in congenital and developmental abnormalities in humans. Importantly, genetic defects in specific metabolic pathways often manifest in tissue-specific patterns indicating the degree of metabolic flexibility is cell type dependent during development, but we currently lack a complete understanding of the capacity for metabolic compensation in the developing embryo and placenta. In this article, we investigate the metabolic resources that support purine metabolism during development by demonstrating the placenta utilizes both the de novo and salvage purine synthesis pathways, and we report the constraint of embryonic use of the salvage pathway during midgestation both under basal conditions or when the de novo pathway is blocked, independent of placental inputs. Our data is consistent with previous reports of embryonic lethality in mouse embryos lacking Impdh2, which cannot be rescued by maternal guanosine supplementation^2^. Our data also provide interesting observations about the regulation of purine metabolism across embryonic tissues and the tight control of hypoxanthine levels during development. Inhibition of Impdh with mizoribine in mouse embryos produces phenotypes similar to what is observed in Lesch-Nyan Syndrome, caused by mutations in HPRT^26,27^, highlighting the cell type specificity of the salvage pathway. Although there have been no reports on the impact of HPRT1 deficiency in the placenta, genetic deficiency of HPRT in induced pluripotent stem cells (iPSCs) alters the development of dopaminergic but not cortical neural progenitor cells although GMP and AMP levels are reduced in both cell types^28^. Interestingly, HPRT deficiency inhibits dopaminergic but not cortical neuron differentiation through decreased GMP levels, degradation of Rheb, and mTORC1 inactivation suggesting that the GMP-mTORC1 metabolic checkpoint may be a common mechanism in some developmental cell types^28^. Embryonic tissues maintain hypoxanthine levels within a tight concentration and our data suggests that excess hypoxanthine in the embryo is used by the placenta to support purine salvage. Finally, we show reduced circulating hypoxanthine levels early in human pregnancy and are further reduced in patients with SGA placentas, which have modestly reduced GMP abundance suggesting that reduced circulating hypoxanthine levels early in pregnancy could be a risk factor for abnormal placental growth.

## Acknowledgments

This work was funded by the University of Texas Southwestern Medical Center, Department of Obstetrics and Gynecology, Cecil H. and Ida Green Center for Reproductive Biology Sciences, and the Howard Hughes Medical Institute. Work was also supported by the NIH/UTSW Nutrition and Obesity Research Center (NIH: P30DK127984). The authors thank vivarium staff Brooke Groff and Crystall Lopez from the Howard Hughes Medical Institute Janelia Research Campus for mouse colony care and timed-pregnant setups. The authors acknowledge the Howard Hughes Medical Institute at Janelia Research Campus Molecular Genomics Shared Resource Core Facility (RRID:SCR_026832), and the Quantitative Genomics Core Facility (RRID:SCR_022694).

## Author contributions

W.X. designed and performed experiments, analyzed data, wrote and edited the manuscript. N.DLC., A.W., and N.K. prepared samples and performed metabolomic LCMS analysis. D.L. established the ex utero culture protocol, a performed embryo culture, immunostainings, imaging. S.W. and M.F-R. collected human serum for ex utero culture experiments. E.L.D., D.B.N., C.Y.S., and C.L.H. provided clinical oversight and guidance, sample availability, and D.D.M., statistics for human studies. J.H.H. supervised executions of experiments and adequate analysis of data. S.S. performed sci-seq3 library preparation and sequencing, and S.G. performed all single cell transcriptomics bioinformatic analyses and assisted on manuscript writing. A.A-C. conceived the idea for this project, established the ex utero culture protocol and gas regulation systems/incubators, designed and conducted most embryology, sequencing, imaging and metabolism experiments, supervised executions of experiments, adequate data analysis, and wrote and edited the manuscript. A.S. conceived the idea for this project, supervised executions of experiments, adequate data analysis, and wrote and edited the manuscript.

---

**Figure S1.**
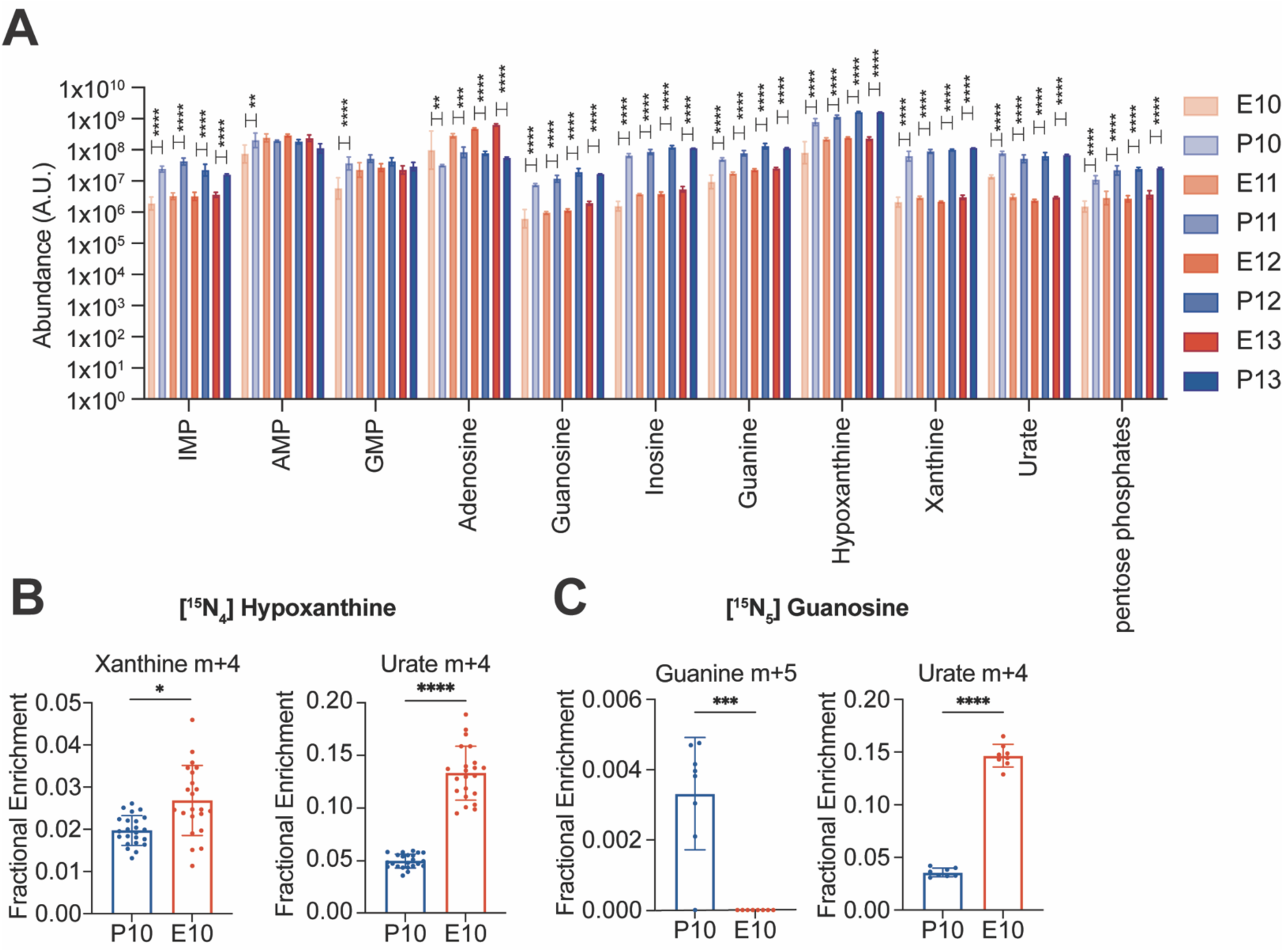
Assessment of salvage purine synthesis pathway in embryo and placenta. **A**. Relative abundance of purines in embryo and placenta on different gestational day (GD10.5-GD13.5). **B**. Fractional enrichment xanthine m+4 and urate m+4 from [^15^N_4_]-Hypoxanthine in embryo(E) and placenta(P). **C**. Fractional enrichment of guanine m+5 and urate m+4 from [^15^N_5_] guanosine in embryo(E) and placenta(P). Statistical tests: (**A**) Two-way ANOVA, Sidak’s multiple comparisons test. (**B-C**) Unpaired, two-tailed t test. ****: P < 0.0001; ***: P < 0.001; **: P < 0.01, *: P < 0.05

**Figure S2.**
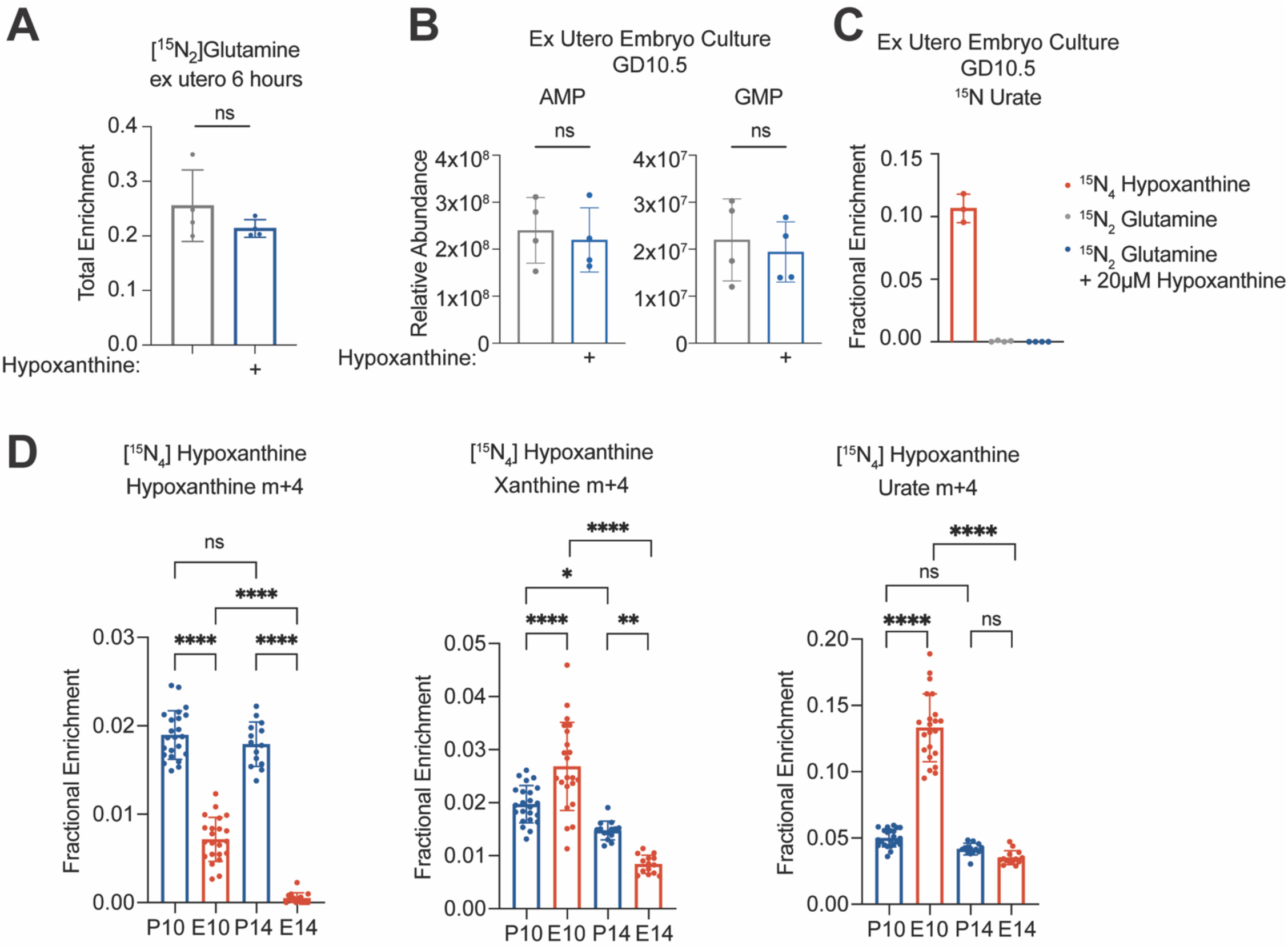
De novo purine synthesis pathway is dominant in embryo. **A**. Fractional enrichment of glutamine from 6h- [^15^N_2_]Glutamine tracing in GD10.5 ex-utero embryo treated with or without 20μM hypoxanthine. **B**. Relative abundance of AMP and GMP in ex utero-embryo treated with or without 20μM hypoxanthine for 6h on GD10.5. **C**. ^15^N labeling enrichment of urate from 6h tracing of [^15^N_4_] hypoxanthine or [^15^N_2_] Glutamine in ex utero embryo. **D**. Fractional enrichment of hypoxanthine m+4, urate m+4 and xanthine m+4 from [^15^N_4_] hypoxanthine infusion in embryo and placenta from GD10.5 and E14.5. Statistical tests: (**A-B**) Unpaired, two-tailed t test. (**D**) (hypoxanthine, xanthine) Welch’s One-way ANOVA tests, Dunnett’s T3 multiple comparisons test; (urate) Ordinary one way ANOVA, Tukey’s multiple comparisons. ****: P < 0.0001; ***: P < 0.001; **: P < 0.01, *: P < 0.05, ns: P>0.05

**Figure S3.**
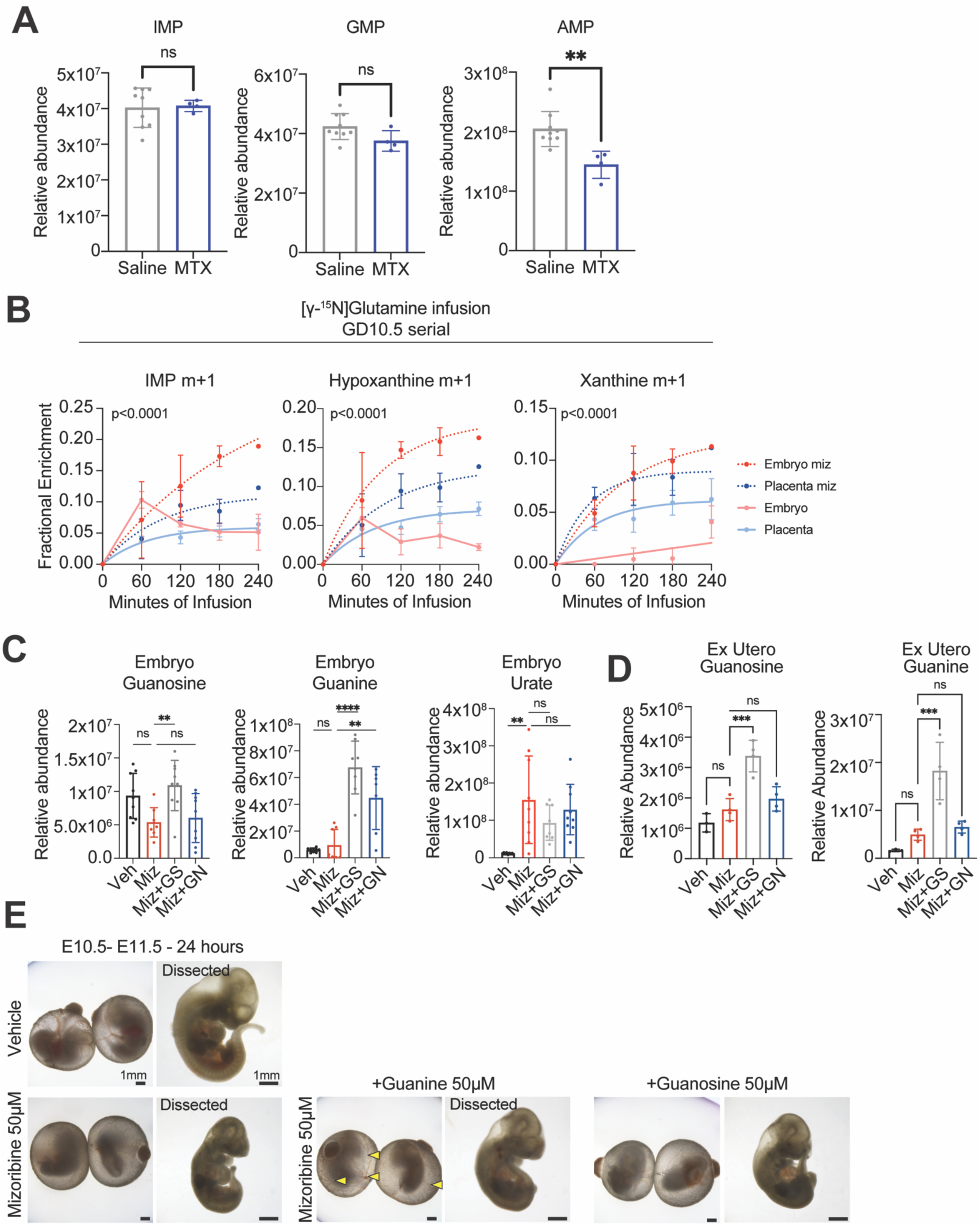
Embryo is incapable of utilizing purine bases for salvage synthesis. **A**. Relative abundance of IMP, GMP and AMP in GD12.5 placenta from mice treated with 5mg/kg MTX from GD10.5-GD12.5. B. Pretreatment (1h) mizoribine(50mg/kg) followed by [γ-^15^N] Glutamine serial caesarian-section, fractional enrichment of IMP m+1, hypoxanthine m+1 and xanthine m+1. **C**. Relative abundance of guanosine, guanine and urate in in utero embryo with indicated treatment from GD10.5-GD12.5. **D**. Relative abundance of guanosine and guanine in ex utero grown embryos with indicated treatment. **E**. Bright field images of ex utero embryos with indicated treatments from GD10.5 to GD11.5. Statistical tests: (**A**) Unpaired, two-tailed t test; (**B**) Plateau followed by one-phase association least-squares fitting, Extra sum-of-squares F test; (**C-D**) (Guanosine, Guanine) One-way ANOVA. Tukey’s multiple comparison tests (Urate) Welch’s one-way ANOVA, Dunnett’s T3 multiple comparisons test. ****: P < 0.0001; ***: P < 0.001; **: P < 0.01, *: P < 0.05, ns: P>0.05

**Figure S4.**
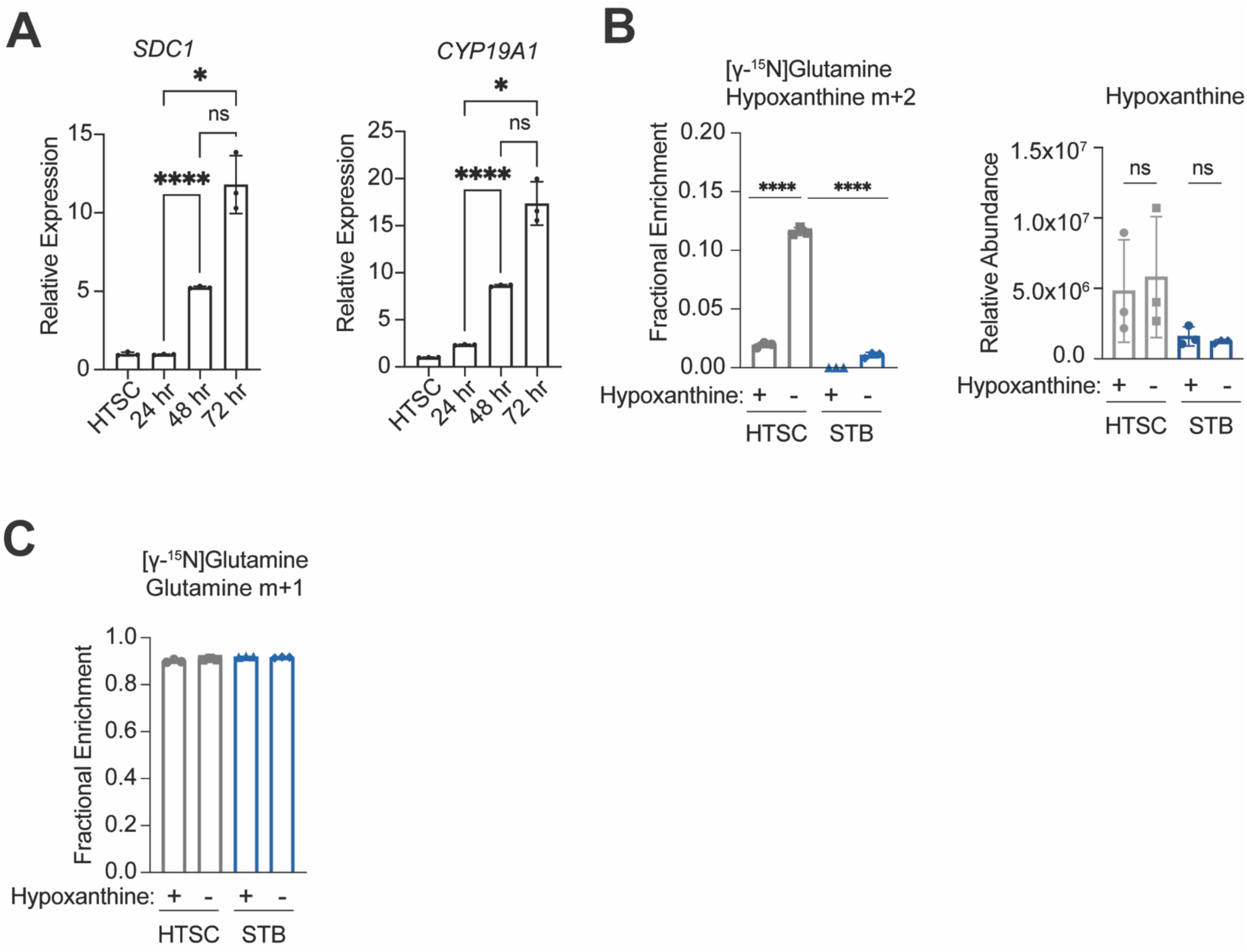
Placenta can utilize purine bases for salvage synthesis. **A**. RT-qPCR analysis of *SDC1* and *CYP19A1* at indicated time points during HTSC differentiation. **B**. Fractional enrichment of hypoxanthine m+2 from [γ-^15^N] Glutamine tracing (left) and relative abundance of hypoxanthine (right) in HTSC and STB cultured in media with or without hypoxanthine. **C**. Fractional enrichment of glutamine m+1 from [γ-^15^N] Glutamine in HTSC and STB cultured in media with or without hypoxanthine. Statistical tests: (**A-B**) Ordinary one-way ANOVA with Tukey’s multiple comparisons. ****: P < 0.0001; ***: P < 0.001; **: P < 0.01, *: P < 0.05, ns: P>0.05

**Figure S5.**
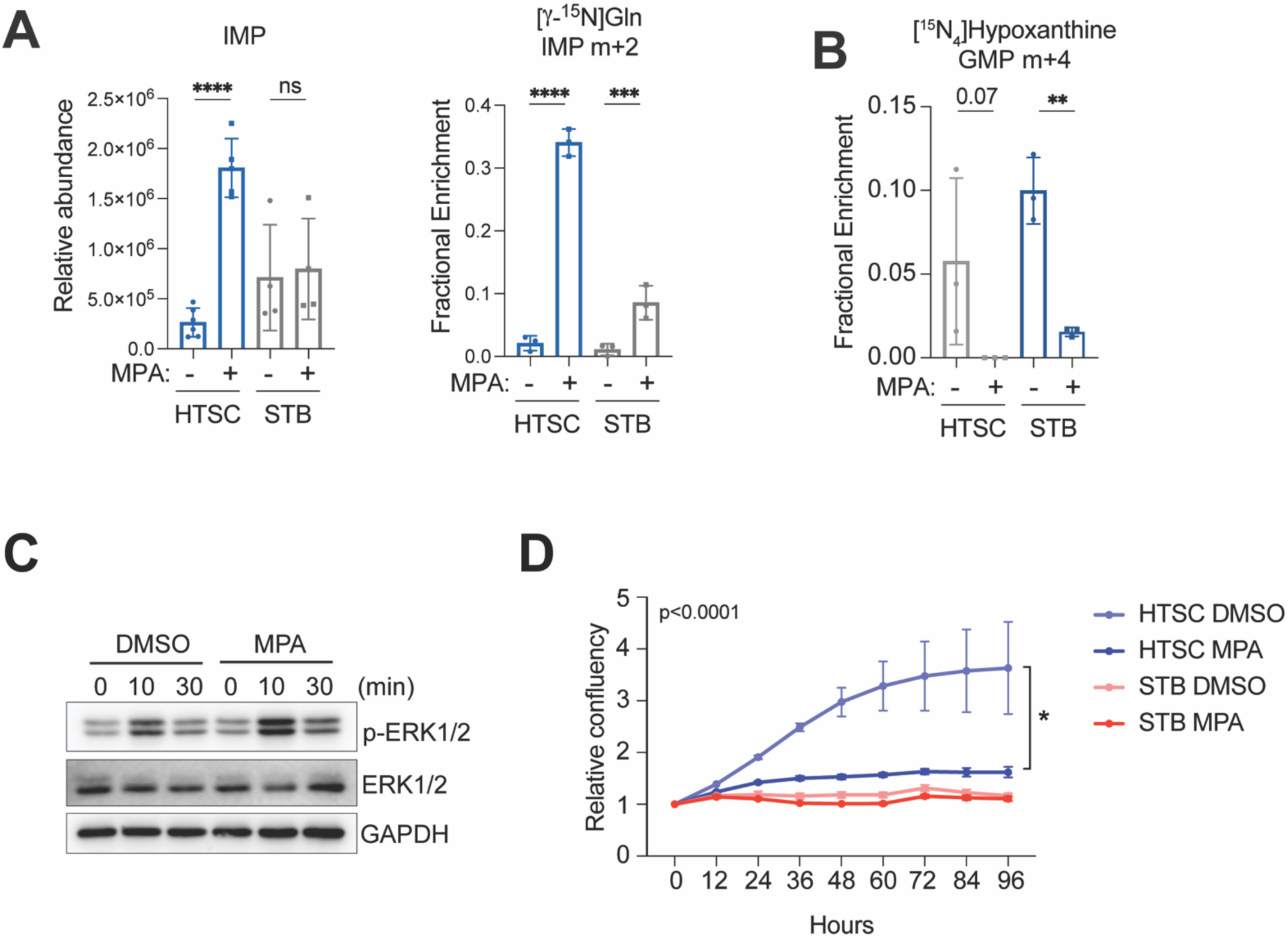
**A**. Relative abundance of IMP (left) and fractional enrichment of IMP m+2 (right) from [γ-^15^N] Glutamine tracing in HTSC and STB after treating with 5 μM MPA for 24h (left). **B**. Fractional enrichment of GMP m+4 from [^15^N_4_] hypoxanthine tracing in HTSC and STB after treating with 5 μM MPA for 24h. **C**. Immunoblots of p-ERK1/2 and ERK1/2 after differentiation for 30min in HTSC with indicated 24h pretreatments. **D**. Proliferation assessment of HTSC or differentiated HTSC with MPA treatment by Incucyte® Live-Cell Analysis System. Statistical tests: **(A-B)** Ordinary One-way ANOVA with Tukey’s multiple comparisons tests; (**D**) Exponential (Malthusian) growth, Extra sum-of-squares F test, Unpaired, two-tailed t tests for 96h time point. ****: P < 0.0001; ***: P < 0.001; **: P < 0.01, *: P < 0.05, ns: P>0.05

## References

1 Gu, J. J. et al. Targeted disruption of the inosine 5’-monophosphate dehydrogenase type I gene in mice. Mol Cell Biol 23, 6702–6712 (2003). 10.1128/MCB.23.18.6702-6712.2003

2 Gu, J. J. et al. Inhibition of T lymphocyte activation in mice heterozygous for loss of the IMPDH II gene. J Clin Invest 106, 599–606 (2000). 10.1172/JCI8669

3 Tran, D. H. et al. De novo and salvage purine synthesis pathways across tissues and tumors. Cell 187, 3602–3618 e3620 (2024). 10.1016/j.cell.2024.05.011

4 Solmonson, A. et al. Compartmentalized metabolism supports midgestation mammalian development. Nature 604, 349–353 (2022). 10.1038/s41586-022-04557-9

5 Tippetts, T. S., Sieber, M. H. & Solmonson, A. Beyond energy and growth: the role of metabolism in developmental signaling, cell behavior and diapause. Development 150 (2023). 10.1242/dev.201610

6 Oginuma, M. et al. Intracellular pH controls WNT downstream of glycolysis in amniote embryos. Nature 584, 98–101 (2020). 10.1038/s41586-020-2428-0

7 Oginuma, M. et al. A Gradient of Glycolytic Activity Coordinates FGF and Wnt Signaling during Elongation of the Body Axis in Amniote Embryos. Dev Cell 40, 342–353 e310 (2017). 10.1016/j.devcel.2017.02.001

8 Cao, D. et al. Selective utilization of glucose metabolism guides mammalian gastrulation. Nature 634, 919–928 (2024). 10.1038/s41586-024-08044-1

9 Galli, L. M., Barnes, T. L., Secrest, S. S., Kadowaki, T. & Burrus, L. W. Porcupine-mediated lipid-modification regulates the activity and distribution of Wnt proteins in the chick neural tube. Development 134, 3339–3348 (2007). 10.1242/dev.02881

10 Lu, C. et al. IDH mutation impairs histone demethylation and results in a block to cell diberentiation. Nature 483, 474–478 (2012). 10.1038/nature10860

11 Zhao, S. et al. ATP-Citrate Lyase Controls a Glucose-to-Acetate Metabolic Switch. Cell Rep 17, 1037–1052 (2016). 10.1016/j.celrep.2016.09.069

12 Longo, L. et al. Maternally transmitted severe glucose 6-phosphate dehydrogenase deficiency is an embryonic lethal. EMBO J 21, 4229–4239 (2002). 10.1093/emboj/cdf426

13 Perez-Garcia, V. et al. Placentation defects are highly prevalent in embryonic lethal mouse mutants. Nature 555, 463–468 (2018). 10.1038/nature26002

14 Aguilera-Castrejon, A. et al. Ex utero mouse embryogenesis from pre-gastrulation to late organogenesis. Nature 593, 119–124 (2021). 10.1038/s41586-021-03416-3

15 Smith, T. K. T. et al. AMPK at the interface of nutrient sensing, metabolic flux and energy homeostasis. Nat Metab 8, 27–51 (2026). 10.1038/s42255-025-01442-3

16 Iriyama, T., Sayama, S. & Osuga, Y. Role of adenosine signaling in preeclampsia. J Obstet Gynaecol Res 48, 49–57 (2022). 10.1111/jog.15066

17 Subiabre, M. et al. Role of insulin, adenosine, and adipokine receptors in the foetoplacental vascular dysfunction in gestational diabetes mellitus. Biochim Biophys Acta Mol Basis Dis 1866, 165370 (2020). 10.1016/j.bbadis.2018.12.021

18 Simmons, D. G. et al. Early patterning of the chorion leads to the trilaminar trophoblast cell structure in the placental labyrinth. Development 135, 2083–2091 (2008). 10.1242/dev.020099

19 Okae, H. et al. Derivation of Human Trophoblast Stem Cells. Cell Stem Cell 22, 50–63 e56 (2018). 10.1016/j.stem.2017.11.004

20 Shpakov, A. O. Hormonal and Allosteric Regulation of the Luteinizing Hormone/Chorionic Gonadotropin Receptor. Front Biosci (Landmark Ed) 29, 313 (2024). 10.31083/j.fbl2909313

21 Ma, L., Chen, Z., Erdjument-Bromage, H., Tempst, P. & Pandolfi, P. P. Phosphorylation and functional inactivation of TSC2 by Erk implications for tuberous sclerosis and cancer pathogenesis. Cell 121, 179–193 (2005). 10.1016/j.cell.2005.02.031

22 Hoxhaj, G. et al. The mTORC1 Signaling Network Senses Changes in Cellular Purine Nucleotide Levels. Cell Rep 21, 1331–1346 (2017). 10.1016/j.celrep.2017.10.029

23 Emmanuel, N. et al. Purine Nucleotide Availability Regulates mTORC1 Activity through the Rheb GTPase. Cell Rep 19, 2665–2680 (2017). 10.1016/j.celrep.2017.05.043

24 Shao, X. et al. Placental trophoblast syncytialization potentiates macropinocytosis via mTOR signaling to adapt to reduced amino acid supply. Proc Natl Acad Sci U S A 118 (2021). 10.1073/pnas.2017092118

25 Pinar, H., Sung, C. J., Oyer, C. E. & Singer, D. B. Reference values for singleton and twin placental weights. Pediatr Pathol Lab Med 16, 901–907 (1996). 10.1080/15513819609168713

26 Witteveen, J. S. et al. HGprt deficiency disrupts dopaminergic circuit development in a genetic mouse model of Lesch-Nyhan disease. Cell Mol Life Sci 79, 341 (2022). 10.1007/s00018-022-04326-x

27 Torres, R. J. & Puig, J. G. Hypoxanthine-guanine phosophoribosyltransferase (HPRT) deficiency: Lesch-Nyhan syndrome. Orphanet J Rare Dis 2, 48 (2007). 10.1186/1750-1172-2-48

28 Bell, S. et al. Lesch-Nyhan disease causes impaired energy metabolism and reduced developmental potential in midbrain dopaminergic cells. Stem Cell Reports 16, 1749–1762 (2021). 10.1016/j.stemcr.2021.06.003

29 Yang, L. et al. Innate immune signaling in trophoblast and decidua organoids defines diberential antiviral defenses at the maternal-fetal interface. Elife 11 (2022). 10.7554/eLife.79794

30 Qiu, C. et al. A single-cell transcriptional timelapse of mouse embryonic development, from gastrula to pup. bioRxiv (2023). 10.1101/2023.04.05.535726

31 Andreatta, M. & Carmona, S. J. UCell: Robust and scalable single-cell gene signature scoring. Comput Struct Biotechnol J 19, 3796–3798 (2021). 10.1016/j.csbj.2021.06.043

